# The Strength of Selection Against Neanderthal Introgression

**DOI:** 10.1101/030148

**Authors:** Ivan Juric, Simon Aeschbacher, Graham Coop

## Abstract

Hybridization between humans and Neanderthals has resulted in a low level of Neanderthal ancestry scattered across the genomes of many modern-day humans. After hybridization, on average, selection appears to have removed Neanderthal alleles from the human population. Quantifying the strength and causes of this selection against Neanderthal ancestry is key to understanding our relationship to Neanderthals and, more broadly, how populations remain distinct after secondary contact. Here, we develop a novel method for estimating the genome-wide average strength of selection and the density of selected sites using estimates of Neanderthal allele frequency along the genomes of modern-day humans. We confirm that East Asians had somewhat higher initial levels of Neanderthal ancestry than Europeans even after accounting for selection. We find that the bulk of purifying selection against Neanderthal ancestry is best understood as acting on many weakly deleterious alleles. We propose that the majority of these alleles were effectively neutral—and segregating at high frequency—in Neanderthals, but became selected against after entering human populations of much larger effective size. While individually of small effect, these alleles potentially imposed a heavy genetic load on the early-generation human–Neanderthal hybrids. This work suggests that differences in effective population size may play a far more important role in shaping levels of introgression than previously thought.

**Author Summary:** A small percentage of Neanderthal DNA is present in the genomes of many contemporary human populations due to hybridization tens of thousands of years ago. Much of this Neanderthal DNA appears to be deleterious in humans, and natural selection is acting to remove it. One hypothesis is that the underlying alleles were not deleterious in Neanderthals, but rather represent genetic incompatibilities that became deleterious only once they were introduced to the human population. If so, reproductive barriers must have evolved rapidly between Neanderthals and humans after their split. Here, we show that oberved patterns of Neanderthal ancestry in modern humans can be explained simply as a consequence of the difference in effective population size between Neanderthals and humans. Specifically, we find that on average, selection against individual Neanderthal alleles is very weak. This is consistent with the idea that Neanderthals over time accumulated many weakly deleterious alleles that in their small population were effectively neutral. However, after introgressing into larger human populations, those alleles became exposed to purifying selection. Thus, rather than being the result of hybrid incompatibilities, differences between human and Neanderthal effective population sizes appear to have played a key role in shaping our present-day shared ancestry.

## Introduction

The recent sequencing of ancient genomic DNA has greatly expanded our knowledge of the relationship to our closest evolutionary cousins, the Neanderthals [1–5]. Neanderthals, along with Denisovans, were a sister group to modern humans, having likely split from modern humans around 550,000–765,000 years ago [5]. Genome-wide evidence suggests that modern humans interbred with Neanderthals after humans spread out of Africa, such that nowadays 1.5–2.1% of the autosomal genome of non-African modern human populations derive from Neanderthals [2]. This admixture dates on average to 47,000–65,000 years ago [6, 51], with potentially a second pulse into the ancestors of populations now present in East Asia [2, 7–10].

While some introgressed archaic alleles appear to have been adaptive in anatomically modern human (AMH) populations [11–13], on average selection has been suggested to act against Neanderthal DNA from modern humans. This can be seen from the non-uniform distribution of Neanderthal alleles along the human genome [8, 12]. In particular, regions of high gene density or low recombination rate have low Neanderthal ancestry, which is consistent with selection removing Neanderthal ancestry more efficiently from these regions [12]. In addition, the X chromosome has lower levels of Neanderthal ancestry and Neanderthal ancestry is absent from the Y chromosome and mitochondria [2, 4, 5, 8, 12, 14, 15]. The genome-wide fraction of Neanderthal introgression in Europeans has recently been shown to have decreased over the past forty thousand years, and, consistent with the action of selection, this decrease is stronger near genes [16]. Finally, a pattern of lower levels of Denisovan ancestry near genes and on the X chromosome in modern humans have also recently been reported [17, 18].

It is less clear why the bulk of Neanderthal alleles would be selected against. Were early-generation hybrids between humans and Neanderthals selected against due to intrinsic genetic incompatibilities? Or was this selection mostly ecological or cultural in nature? If reproductive barriers had already begun to evolve between Neanderthals and AMH, then these two hominids may have been on their way to becoming separate species before they met again [12, 19, 20]. Or, as we propose here, did differences in effective population size and resulting genetic load between humans and Neanderthals shape levels of Neanderthal admixture along the genome?

We set out to estimate the average strength of selection against Neanderthal alleles in AMH. Due to the relatively short divergence time of Neanderthals and AMH, we still share much of our genetic variation with Neanderthals. However, we can recognize alleles of Neanderthal ancestry in humans by aggregating information along the genome using statistical methods [8, 12]. Here, we develop theory to predict the frequency of Neanderthal-derived alleles as a function of the strength of purifying selection at linked exonic sites, recombination, initial introgression proportion, and split time. We fit these predictions to recently published estimates of the frequency of Neanderthal ancestry in modern humans [12]. Our results enhance our understanding of how selection shaped the genomic contribution of Neanderthal to our genomes, and shed light on the nature of Neanderthal–human hybridization.

## Results

In practice, we do not know the location of the deleterious Neanderthal alleles along the genome, nor could we hope to identify them all as some of their effects may be weak (but perhaps important in aggregate). Therefore, we average over the uncertainty in the locations of these alleles. We assume that each exonic base independently harbors a deleterious Neanderthal allele with probability *μ*. Building on a long-standing theory on genetic barriers to gene flow [21–23, 25, 26], at each neutral site ℓ in the genome, we can express the present-day expected frequency of Neanderthal alleles in our admixture model in terms of the initial frequency *p*_0_, as well as a function *g_ℓ_* of the recombination rates **r** between ℓ and the neighboring exonic sites under selection, and the parameters *s*, *t*, and *μ* (see Eq. 5, S2 Text). That is, at locus ℓ, a fraction *p*_ℓ, *t*_ = *p*_0_*g*_ℓ_(**r**, *s, t, μ*) of modern humans are expected to carry the Neanderthal allele. The function *g_ℓ_*() decreases with tighter linkage to potentially deleterious sites, larger selection coefficient (*s*), longer time since admixture (*t*), and higher density of deleterious exonic sites (*μ*). If a neutral Neanderthal allele is initially *completely* unassociated with deleterious alleles, *p*_*ℓ,t*_ would on average be equal to *p*_0_. Our model explicitly accounts only for deleterious alleles that are physically linked to a neutral allele. However, in practice, neutral Neanderthal alleles will initially be associated (i.e. in linkage disequilibrium) not only with some linked, but also with potentially many unlinked deleterious alleles. This is because *F*_1_ hybrids inherited half of their genome from Neanderthal parents, which leads to a statistical association even among unlinked Neanderthal-derived alleles. Therefore, *p*_0_ should be thought of as an *effective* initial admixture proportion in the sense that it implicitly absorbs the effect of these physically unlinked, but statistically associated deleterious Neanderthal allele. Technically this is because the effect of unlinked loci (assuming multiplicative fitness) can be factored into a constant multiplier of *g*_ℓ_(), and so can be accomodated into the model by rescaling *p*_0_ (see pages 35 and 36 of [22]) In practice, this means that our estimates of *p*_0_ will almost certainly be underestimating the actual proportion of Neanderthal admixture. We will return to this point in the Discussion. We emphasize that, independently of the effect of unlinked deleterious mutations, there may still be more than one linked deleterious mutation associated with any given focal neutral site on average. To assess this possibility, in S2 Text we compare models that explicitly account for one versus multiple linked deleterious mutations.

To estimate the parameters of our model (*p*_0_, *s*, and *μ*), we minimised the residual sum of squared deviations (RSS) between observed frequencies of Neanderthal alleles [12] and those predicted by our model (see Eq. 6 and S2 Text). We assess the uncertainty in our estimates by bootstrapping large contiguous genomic blocks and re-estimating our parameters. We then provide block-wise bootstrap confidence intervals (CI) based on these (Methods and S2 Text). In Fig 2 and 3, we show the RSS surfaces for the parameters *p*_0_, *s*, and *μ* for autosomal variation in Neanderthal ancestry in the EUR and ASN populations.

**Figure 1.**
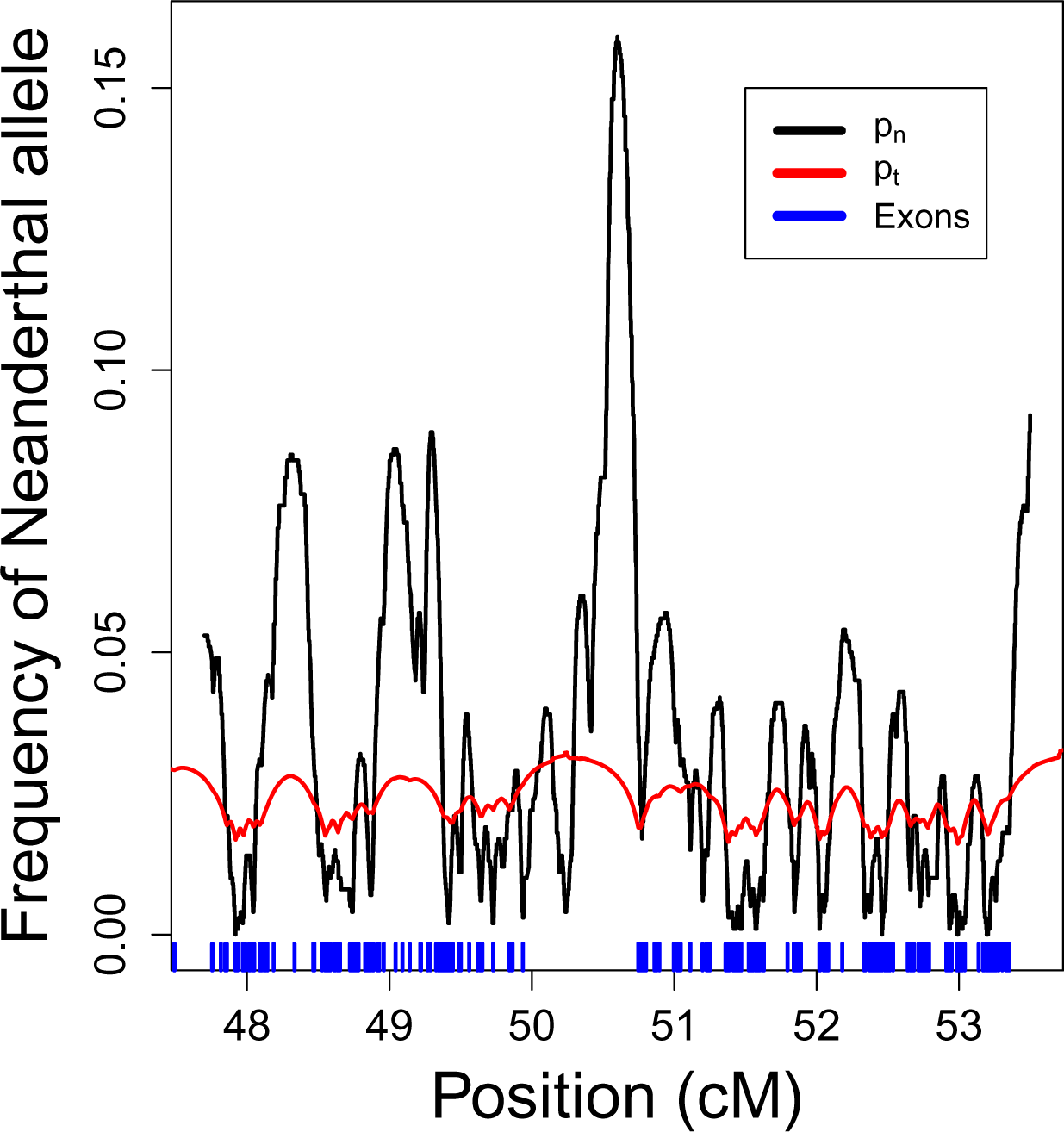
A section of chromosome 1 showing the estimated Neanderthal frequency (*p_n_*, black line) for the EUR sample from [12] and the expected frequency (*p_t_*, red line) predicted by our best fitting model. The midpoints of exons are shown as blue bars. Note that the estimated frequency is expected to have much greater variance along the genome than our prediction due to genetic drift. Our prediction refers to the mean around which the deviation due to genetic drift is centered (S2 Text).

**Figure 2.**
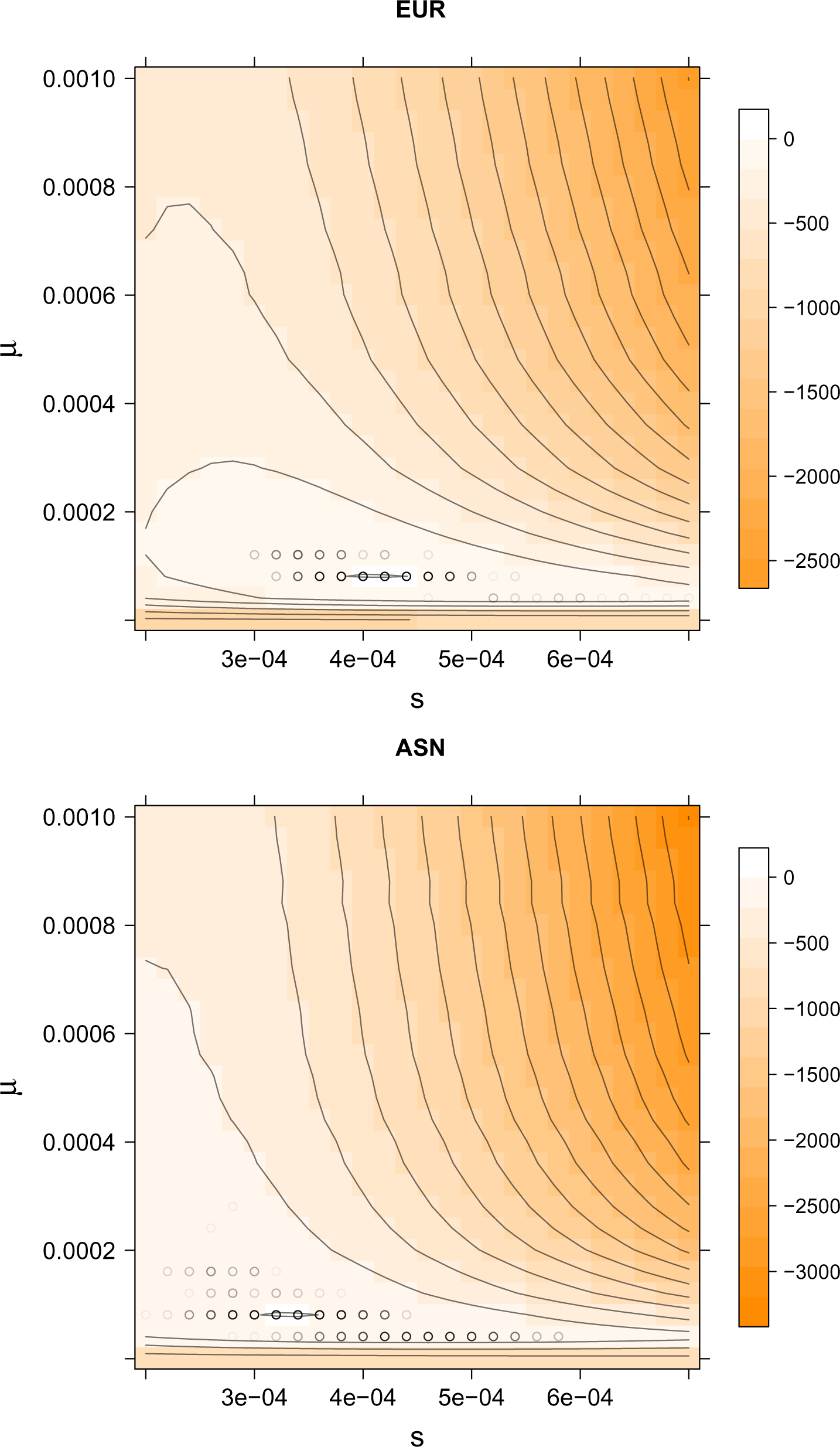
The scaled RSS surface (RSS_min_ - RSS) as a function of *s* and *μ*for EUR and ASN autosomal chromosomes. Each value of the RSS is minimized over *p*_0_, making this a profile RSS surface. Regions in darker shades of orange represent parameter values of lower scaled RSS. Black circles show bootstrap results of 1000 blockwise bootstrap reestimates, with darker circles corresponding to more common bootstrap estimates.

**Figure 3.**
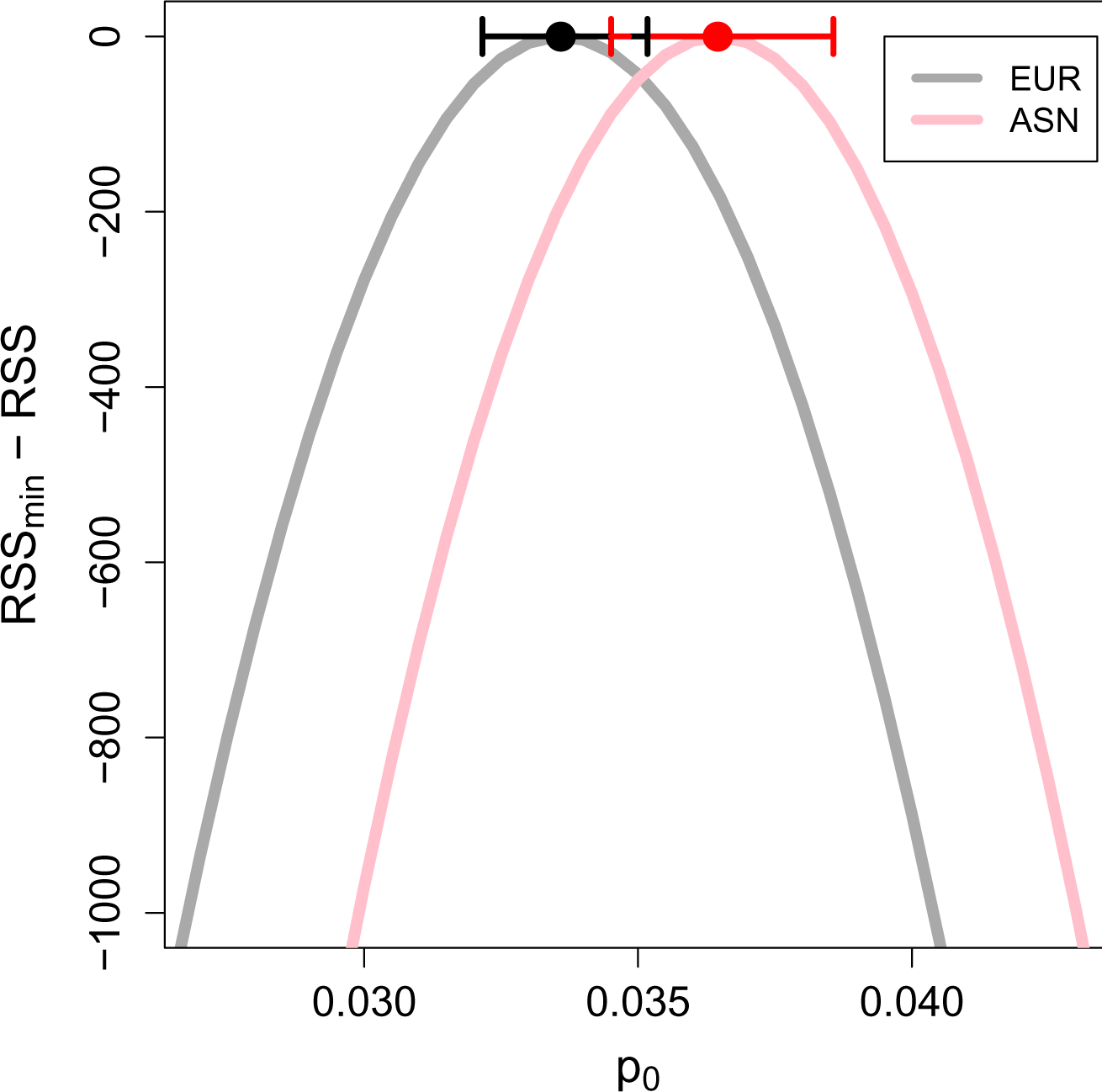
The scaled RSS surface (RSS_min_ - RSS) of autosomal chromosomes as a function of the initial admixture proportion *p*_0_. Results are shown for a model where only the nearest-neighboring exonic site under selection is considered, and for *t* = 2000 generations after the Neanderthal admixture event into the ancestors of EUR (grey) and ASN (pink) populations. Dots and horizontal lines show the value of *p*_0_ that minimizes the RSS and the respective 95% block-bootstrap confidence intervals. The RSS surfaces are shown for values of the selection coefficient (*s*) and exonic density of selection (*μ*) given in Table 1.

For autosomal chromosomes, our best estimates for the average strength of selection against deleterious Neanderthal alleles are low in both EUR and ASN (Fig 2), but statistically different from zero (*s*_EUR_ = 4.1 *×* 10^−4^; 95% CI [3.4 *×* 10^−4^, 5.2 *×* 10^−4^], *s*_ASN_ = 3.5 *×* 10^−4^; 95% CI [2.6 *×* 10^−4^, 5.4 *×* 10^−4^]). We obtain similar estimates if we assume that the Neanderthal ancestry in humans has reached its equilibrium frequency or if we account for the effect of multiple selected sites (see S2 Text). However, and as expected, the estimated selection coefficients are somewhat lower for those models (S2 Text Table S1). Our estimates of the probability of any given exonic site being under selection are similar and low for both samples (*μ*_*EUR*_ = 8.1 *×* 10^−5^; 95% CI [4.1 *×* 10^−5^, 1.2 *×* 10^−4^], *μ*_*ASN*_ = 6.9 *×* 10^−5^; 95% CI [4.1 *×* 10^−5^, 1.6 *×* 10^−4^]). These estimates correspond to less than 1 in 10000 exonic base pairs harboring a deleterious Neanderthal allele, on average. As a result, our estimates of the average selection coefficient against an exonic base pair (the compound parameter (*μs*) are very low, on the order of 10^−8^ in both samples (Table 1).

**Table 1.** Point estimates and 95% bootstrap confidence intervals for the focal parameters. Estimates are based on a minimization of the residual sum of squared deviations (RSS) between observations and a model in which, for each neutral site, only the nearest-neighboring exonic site under selection is considered. Introgression is assumed to have happened *t* = 2000 generations ago.

| Sample | Chr. | $p_0$ | $s \times 10^{-4}$ | $\mu \times 10^{-4}$ | $\mu s \times 10^{-8}$ |
| --- | --- | --- | --- | --- | --- |
| EUR | Auto. | 0.0338 [0.0322, 0.0352] | 4.12 [3.4, 5.2] | 0.81 [0.41, 1.2] | 3.38 [2.59, 4.38] |
| EUR | X | 0.0292 [0.0232, 0.0353] | 9.60 [6.4, 20.8] | 0.81 [0.41, 1.6] | 7.78 [3.28, 15.4] |
| ASN | Auto. | 0.0360 [0.0345, 0.0386] | 3.52 [2.6, 5.4] | 0.69 [0.41, 1.6] | 2.43 [1.48, 4.19] |
| ASN | X | 0.0298 [0.0236, 0.039] | 1.6 [0, 40] | 6.8 [0.01, 10] | 10.88 [0, 32.6] |

Consistent with previous findings [9, 10], we infer a higher initial frequency of Neanderthal alleles in the East Asian sample compared to the European sample (*p*_0, *EUR*_ = 3.38 *×* 10^−2^; 95% CI [3.22 *×* 10^−2^, 3.52 *×* 10^−2^], *p*_0, *ASN*_ = 3.60 *×* 10^−2^; 95% CI [3.45 *×* 10^−2^, 3.86 *×* 10^−2^]), but the 95% bootstrap CI overlap (Fig 3). This occurs because our estimates of the initial frequency of Neanderthal alleles (*p*_0_) are mildly confounded with estimates of the strength of selection per exonic base (*μs*). That is, somewhat similar values of the expected present-day Neanderthal allele frequency can be inferred by simultaneously reducing *p*_0_ and *μs* (Fig 4). This explains why the marginal confidence intervals for *p*_0_ overlap for ASN and EUR. However, if *μs*, the fitness cost of Neanderthal introgression per exonic base pair, is the same for ASN and EUR (i.e. if we take a vertical slice in Fig 4), the values of *p*_0_ for the two samples do not overlap.

**Figure 4.**
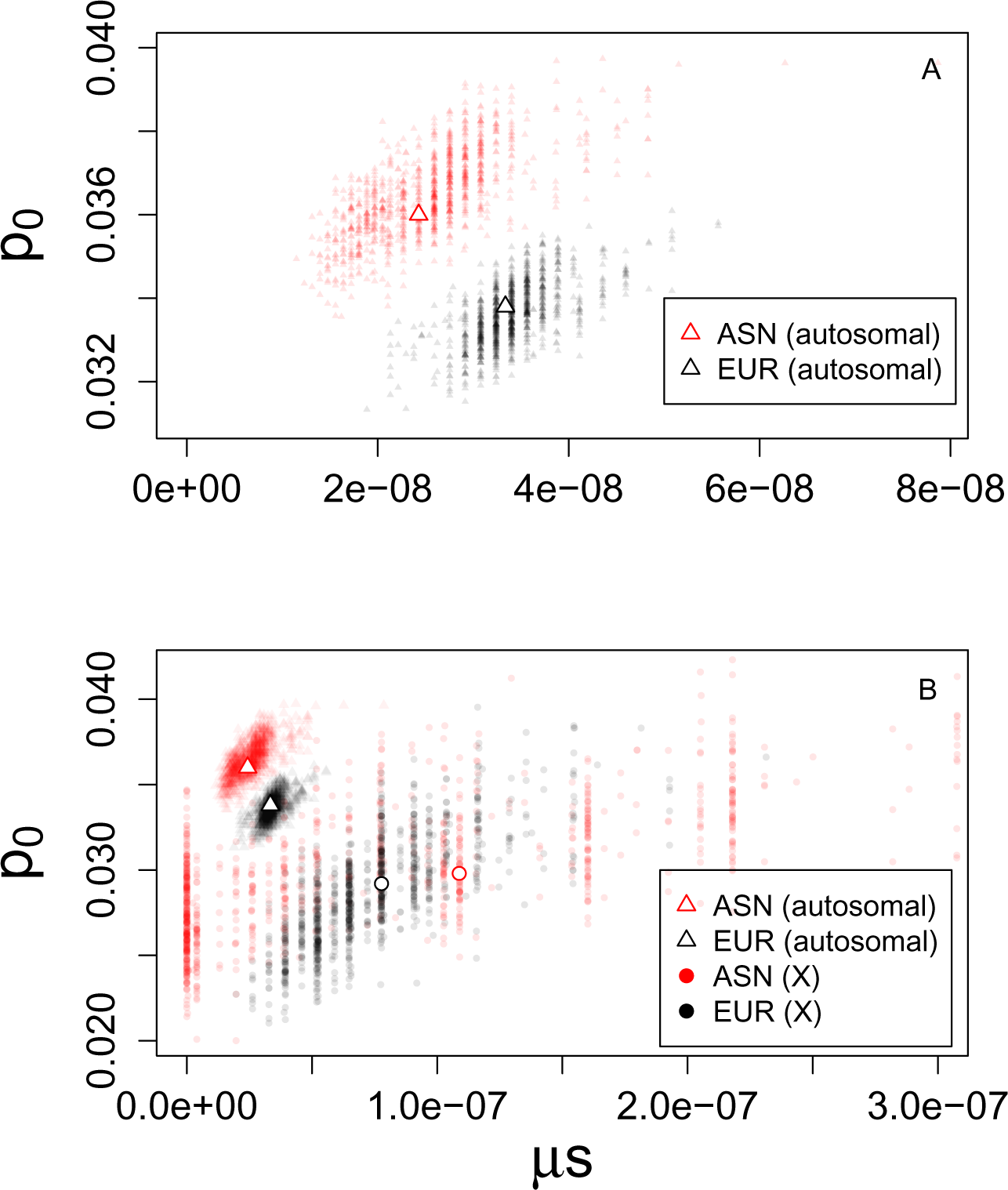
The contrast between the inferred parameters for the East Asian (ASN) and European (EUR) samples for the autosomes (A) and both the X and the autosomes (B). Plots show bootstrap estimates of the initial admixture proportion *p*_0_ against the estimated exonic density of selection *μs*, with the empty symbols denoting our minimum RSS estimates. The clear separation of the point clouds for autosomes and the X for both EUR and ASN modern humans suggests that the combination of selection and initial admixture level are likely the reason why the present-day frequency of Neanderthal alleles differs between autosomal and X chromosomes. Note the different scales of the axes in panels A and B.

To verify the fit of our model, we plot the average observed frequency of Neanderthal alleles, binned by gene density per map unit, and compare it to the allele frequency predicted by our model based on the estimated parameter values (Fig 6). There is good agreement between the two, suggesting that our model provides a good description of the relationship between functional density, recombination rates, and levels of Neanderthal introgression. At the scale of 1 cM, the Pearson correlation between observed and predicted levels of autosomal Neanderthal introgression is 0.897 for EUR and 0.710 for ASN (see Table S3 for a range of other scales).

Our estimated coefficients of selection (*s*) against deleterious Neanderthal alleles are very low, on the order of the reciprocal of the effective population size of humans. This raises the intriguing possibility that our results are detecting differences in the efficacy of selection between AMH and Neanderthals. Levels of genetic diversity within Neanderthals are consistent with a very low long-term effective population size compared to AMH, i.e. a higher rate of genetic drift [5]. This suggests that weakly deleterious exonic alleles may have been effectively neutral and drifted up in frequency in Neanderthals [27–29], only to be slowly selected against after introgressing into modern human populations of larger effective size. To test this hypothesis, we simulated a simple model of a population split between AMH and Neanderthals, using a range of plausible Neanderthal population sizes after the split. In these simulations, the selection coefficients of mutations at exonic sites are drawn from an empirically supported distribution of fitness effects [30]. We track the frequency of deleterious alleles at exonic sites in both AMH and Neanderthals, and compare these frequencies at the time of secondary contact (admixture). We show a subset of our simulation results in Fig 5. Due to a lower effective population size, the simulated Neanderthal population shows an excess of fixed deleterious alleles compared to the larger human population (Fig 5A). This supports the assumption we made in our inference procedure that the deleterious introgressing alleles had been fixed in Neanderthals prior to admixture. Moreover, our estimates of *s* fall in a region of parameter space for which simulations suggest that Neanderthals have a strong excess of population-specific fixed deleterious alleles, compared to humans (Fig 5B). Over the relevant range of selection coefficients, the fraction of simulated exonic sites that harbor these Neanderthal-specific weakly deleterious alleles is on the order of 10^−5^, which is in approximate agreement with our estimates of *μ*. Therefore, a model in which the bulk of Neanderthal alleles, which are now deleterious in modern humans, simply drifted up in frequency due to the smaller effective population size of Neanderthals seems quite plausible. This conclusion has also been independently reached by a recent study via a simulation-based approach [31].

**Figure 5.**
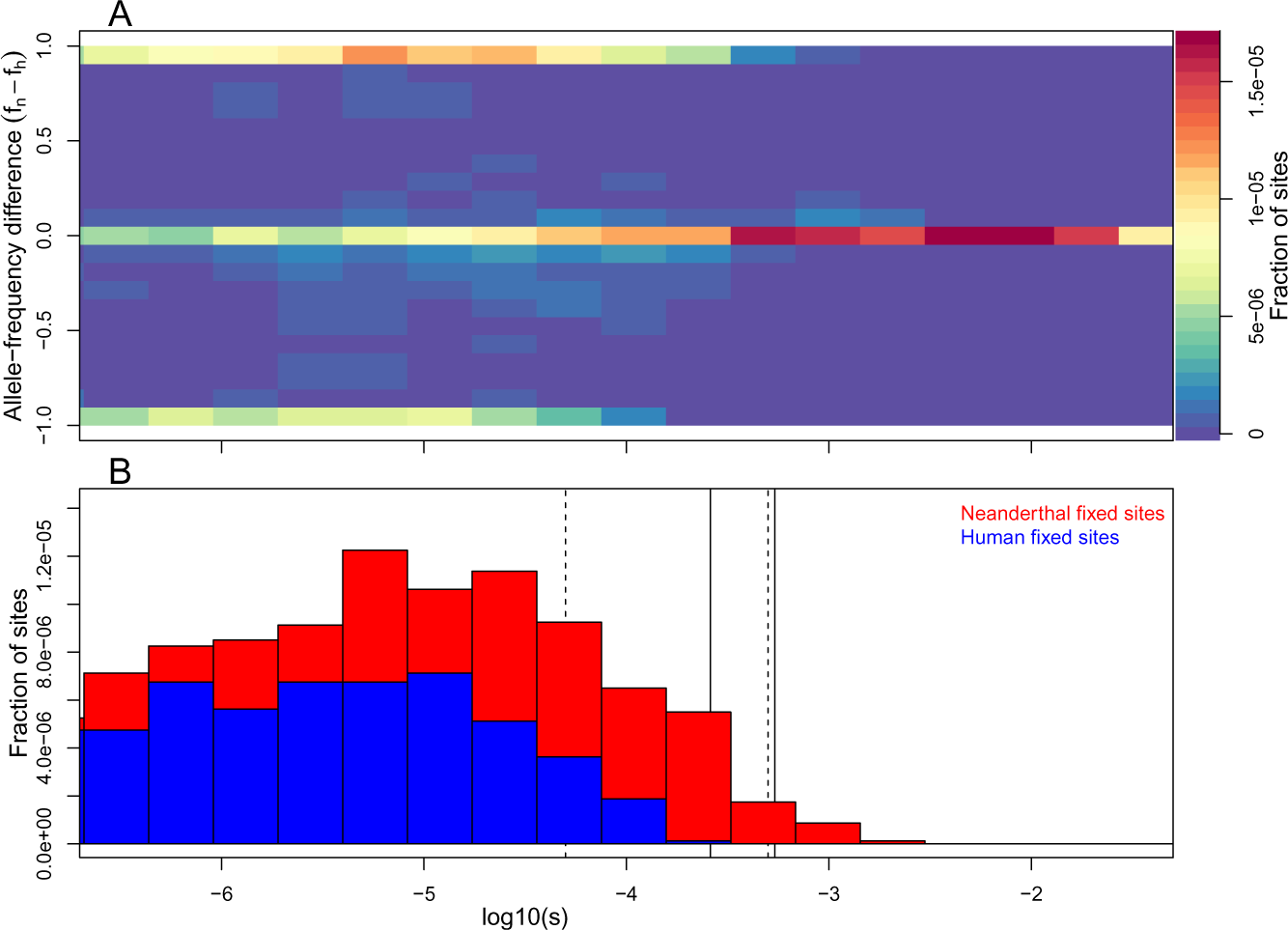
Simulations showing that the Neanderthal population is predicted to harbor an excess of weakly deleterious fixed alleles compared to humans. (A) A two-dimensional histogram of the difference in allele frequency between Neanderthal and human population, and the deleterious selection coefficient over all simulated sites. (B) The fraction of sites in the simulations where there is a human- or Neanderthal-specific fixed difference, binned by selection coefficient. Dotted lines indicate the nearly-neutral selection coefficient (i.e. the inverse of the effective population size) for Neanderthal (right) and Human (left) populations. Solid lines show the 95% CI of *s* for ASN (the larger of the two CI) that we inferred. Note that monomorphic sites are not shown, but are included in the denominator of the fraction of sites.

**Figure 6.**
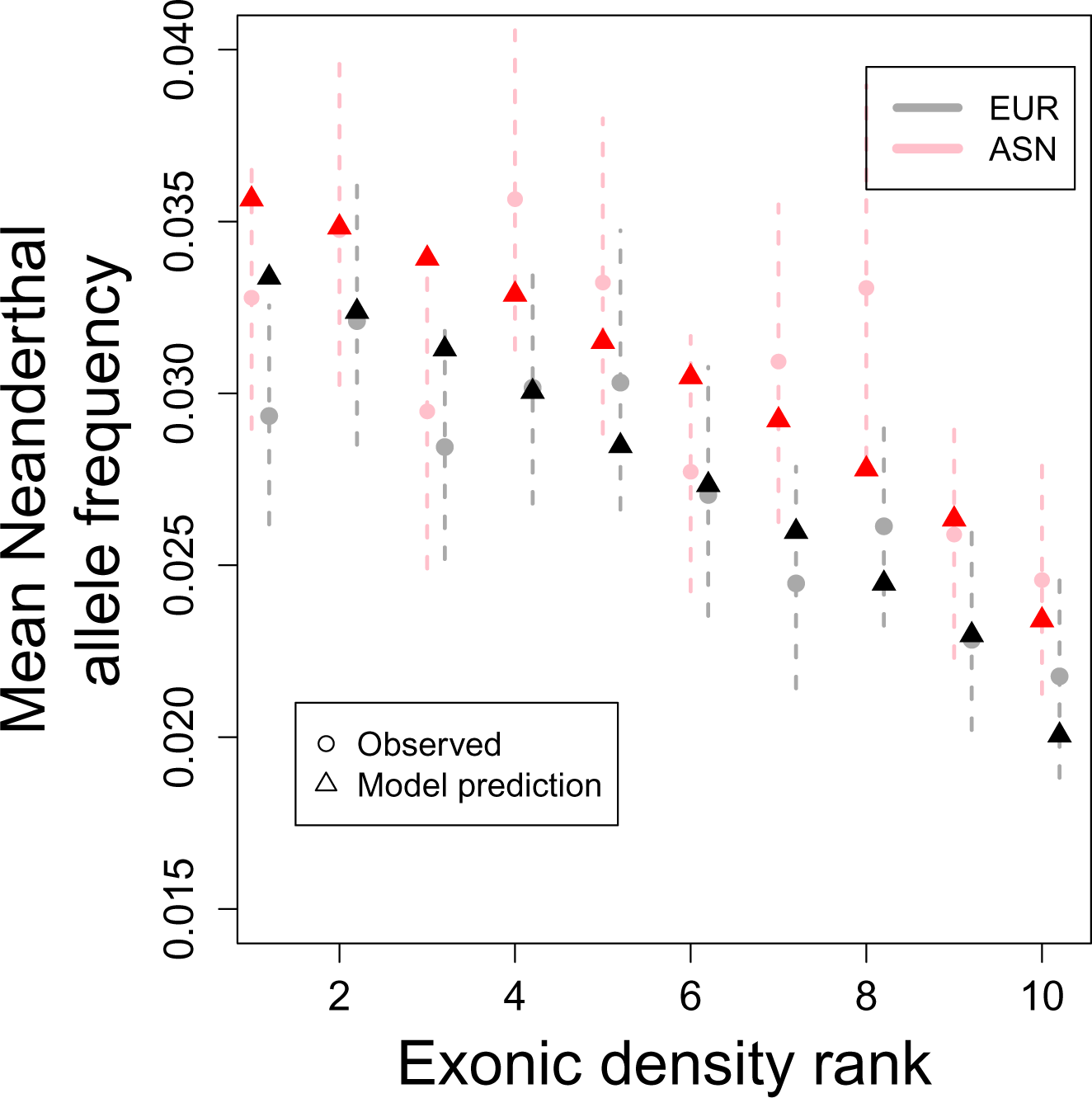
Genomic regions with lower exonic density contain higher average Neanderthal allele frequency in both in Europeans (grey circles) and Asians (pink circles). We find a good fit to this pattern under our model (black and red triangles). Ranks are obtained by splitting the genome into 1 cM segments, calculating the number of exonic sites for each segment and sorting the segments into ten bins of equal size. Dashed lines represent 95% blockwise bootstrap confidence intervals. Plots created for different segment sizes look similar (S2 Text).

We finally turn to the X chromosome, where observed levels of Neanderthal ancestry are strongly reduced compared to autosomes [8, 12]. This reduction could be consistent with the X chromosome playing an important role in the evolution of hybrid incompatibilities at the early stages of speciation [12]. However, a range of other phenomena could explain the observed difference between the X and autosomes, including sex-biased hybridization among populations, the absence of recombination in males, as well as differences in the selective regimes [32–34]. We modified our model to reflect the transmission rules of the X chromosome and the absence of recombination in males. We give the X chromosome its own initial level of introgression (*p*_0, *X*_), different from the autosomes, which allows us to detect a sex bias in the direction of matings between AHM and Neanderthals. Although our formulae can easily incorporate sex-specific selection coefficients, we keep a single selection coefficient (*s_X_*) to reduce the number of parameters. Therefore, *s_X_* reflects the average reduction in relative fitness of deleterious Neanderthal alleles across heterozygous females and hemizygous males.

We fit the parameters *p*_0_, *X*, *μ*_*X*_, and *s_X_* using our modified model to [12]’s observed levels of admixture on the X chromosome (Table 1; S12 Fig and S13 Fig). Given the smaller amount of data, the inference is more challenging as the parameters are more strongly confounded (for example of *μ*_*X*_ and *s_X_*, see S12 Fig and S13 Fig). We therefore focus on the compound parameter *μ*_*X*_*s*_*X*_, i.e. the average selection coefficient against an exonic base pair on the X. In Fig 4, we plot a sample of a thousand bootstrap estimates of *μ*_*X*_*s*_*X*_ for the X, along with analogous estimates of *μs* for autosomal chromosomes. For the X chromosome, there is also strong confounding between *p*_0, *X*_ and *μ*_*X*_*s*_*X*_, to a much greater extent than on the autosomes (note the larger spread of the X point clouds). Due to this confounding, our marginal confidence intervals for *μ*_*X*_*s*_*X*_ and *p*_0, *X*_ overlap with their autosomal counterparts (Table 1). However, the plot of *p*_0_ and *μs* bootstrap estimates clearly shows that the X chromosome and autosomes differ in their parameters.

For reasons we do not fully understand, the range of parameter estimates for the X chromosome with strong bootstrap support is much larger for the ASN than for the EUR samples (Fig 4). For the ASN samples, the confidence intervals for *μ*_*X*_*s*_*X*_ include zero, suggesting there is no strong evidence for selection against introgression on the X. This is consistent with the results of [12], who found only a weakly significant correlation between the frequency of Neanderthal alleles and gene density on the X chromosome. However, as the ASN confidence intervals for *μ*_*X*_*s*_*X*_ are large and also overlap with the autosomal estimates, it is difficult to say if selection was stronger or weaker on the X chromosome compared to the autosomes. For the EUR samples, however, the confidence intervals for *μ*_X_*s*_*X*_ do not include zero, which suggests significant evidence for selection against introgression on the X, potentially stronger than that on the autosomes. Note that the selection coefficients on the X (*s_X_*, Table 1) are still on the order of one over the effective population size of modern humans, as was the case for the autosomes. Therefore, differences in effective population size between Neanderthals and modern humans, and hence in the efficacy of selection, might well explain observed patterns of introgression on the X as well as on the autosomes. If the exonic density of selection against Neanderthal introgression was indeed stronger on the X, one plausible explanation is the fact that weakly deleterious alleles that are partially recessive would be hidden from selection on the autosomes but revealed on the X in males [32–34].

Our results are potentially consistent with the notion that the present-day admixture proportion on the X chromosome was influenced not only by stronger purifying selection, but also by a lower initial admixture proportion *p*_0, *X*_ (Fig 4). Lower *p*_0, *X*_ is consistent with a bias towards matings between Neanderthal males and human females, as compared to the opposite. Based on our point estimates, and if we attribute the difference between the initial admixture frequency between the X and the autosomes (*p*_0, *X*_ and *p*_0*,A*_) exclusively to sex-biased hybridization, our result would imply that matings between Neanderthal males and human females were about three times more common than the opposite pairing (S2 Text). However, as mentioned above, there is a high level of uncertainty about our X chromosome point estimates. Therefore, we view this finding as very provisional.

## Discussion

There is growing evidence that selection has on average acted against autosomal Neanderthal alleles in anatomically modern humans (AMH). Our approach represents one of the first attempts to estimate the strength of genome-wide selection against introgression between populations. The method we use is inspired by previous efforts to infer the strength of background selection and selective sweeps from their footprint on linked neutral variation on a genomic scale [35–38]. We have also developed an approach to estimate selection against on-going maladaptive gene flow using diversity within and among populations (Aeschbacher and Coop, in prep.) that will be useful in extending these findings to a range of taxa. Building on these approaches, more refined models of selection against Neanderthal introgression could be developed. These could extend our results by estimating a distribution of selective effects against Neanderthal alleles, or by estimating parameters separately for various categories of sequence, such as non-coding DNA, functional genes, and other types of polymorphism(e.g. structural variation) [39].

Here, we have shown that observed patterns of Neanderthal ancestry in modern human populations are consistent with genome-wide purifying selection against many weakly deleterious alleles. For simplicity, we allowed selection to act only on exonic sites. It is therefore likely that the effects of nearby functional non-coding regions are subsumed in our estimates of the density (*μ*) and average strength (*s*) of purifying selection. Therefore, our findings of weak selection are conservative in the sense that the true strength of selection per base pair may be even weaker. We argue that the bulk of selection against Neanderthal ancestry in humans may be best understood as being due to the accumulation of alleles that were effectively neutral in the Neanderthal population, which was of relatively small effective size. However, these alleles started to be purged, by weak purifying selection, after introgressing into the human population, due to its larger effective population size.

Thus, we have shown that it is not necessary to hypothesize many loci harboring intrinsic hybrid incompatibilities, or alleles involved in ecological differences, to explain the bulk of observed patterns of Neanderthal ancestry in AMH. Indeed, given a rather short divergence time between Neanderthals and AMH, it is *a priori* unlikely that strong hybrid incompatibilities had evolved at a large number of loci before the populations interbred. It often takes millions of years for hybrid incompatibilities to evolve in mammals [40, 41], although there are exceptions to this [42], and theoretical results suggest that such incompatibilities are expected to accumulate only slowly at first [43, 44]. While this is a subjective question, our results suggest that genomic data—although clearly showing a signal of selection against introgression—do not strongly support the view that Neanderthals and humans should be viewed as incipient species. Sankararaman *et al.* [12] found that genes expressed in the human testes showed a significant reduction in Neanderthal introgression, and interpreted this as being potentially consistent with a role of reproductive genes in speciation. However, this pattern could also be explained if testes genes were more likely to harbor weakly deleterious alleles, which could have accumulated in Neanderthals. These two hypotheses could be addressed by relating within-species estimates of the distribution of selective effects with estimates of selection against introgression at these testes genes.

This is not to say that alleles of larger effect, in particular those underlying ecological or behavioral differences, did not exist, but rather that they are not needed to explain the observed relationship between gene density and Neanderthal ancestry. Alleles of large negative effect would have quickly been removed from admixed populations, and would likely have led to extended genomic regions showing a deficit of Neanderthal ancestry as described by [8, 12, 45]. Since our method allows us to model the expected amount of Neanderthal ancestry along the genome accounting for selection, it could serve as a better null model for finding regions that are unusually devoid of Neanderthal ancestry.

We have ignored the possibility of adaptive introgressions from Neanderthals into humans. While a number of fascinating putatively adaptive introgressions have come to light [13], and more will doubtlessly be identified, they will likely make up a tiny fraction of all Neanderthal haplotypes. We therefore think that they can be safely ignored when assessing the long-term deleterious consequences of introgression.

As our results imply, selection against deleterious Neanderthal alleles was very weak on average, such that, after tens of thousands of years since their introduction, these alleles will have only decreased in frequency by 56% on average. Thus, roughly seven thousand loci (*≈ μ ×* 82 million exonic sites) still segregate for deleterious alleles introduced into Eurasian populations via interbreeding with Neanderthals. However, given that the initial frequency of the admixture was very low, we predict that a typical EUR or ASN individual today only carries roughly a hundred of these weak-effect alleles, which may have some impact on genetic load within these populations.

Although selection against each deleterious Neanderthal allele is weak, the early-generation human–Neanderthal hybrids might have suffered a substantial genetic load due to the sheer number of such alleles. The cumulative contribution to fitness of many weakly deleterious alleles strongly depends on the form of fitness interaction among them, but we can still make some educated guesses (the caveats of which we discuss below). If, for instance, the interaction was multiplicative, then an average F1 individual would have experienced a reduction in fitness of 1 − (1 − 4 × 10^−4^)^7000^ ≈ 94% compared to modern humans, who lack all but roughly one hundred of these deleterious alleles. This would obviously imply a substantial reduction in fitness, which might even have been increased by a small number of deleterious mutations of larger effect that we have failed to capture. This potentially substantial genetic load has strong implications for the interpretation of our estimate of the effective initial admixture proportion (*p*_0_), and, more broadly, for our understanding of those early hybrids and the Neanderthal population. We now discuss these topics in turn.

Strictly, under our model, the estimate of *p*_0_ reflects the initial admixture proportion in the absence of unlinked selected alleles. However, the large number of deleterious unlinked alleles present in the first generations after admixture violates that assumption, as each of these unlinked alleles also reduces the fitness of hybrids [22]. These unlinked deleterious alleles should cause a potentially rapid initial loss of Neanderthal ancesty following the hybridization. Harris and Nielsen [31] have recently independently conducted simulations of the dynamics of deleterious alleles during the initial period following Neanderthal admixture. They have shown that the frequency of Neanderthal-derived alleles indeed decreases rapidly in the initial generations due to the aggregate effects of many weakly deleterious loci. The reduction in neutral Neanderthal ancestry due to unlinked sites under selection is felt equally along the genome and as such, our estimate of *p*_0_ is an effective admixture proportion that incorporates the genome-wide effect of unlinked deleterious mutations, but not the localized effect of linked deleterious mutations (as formalized by Bengtsson [22]). In practice, segregation and recombination during meiosis in the early generations after admixture will have led to a rapid dissipation of the initial associations (statistical linkage disequilibrium) among any focal neutral site and unlinked deleterious alleles. Therefore, our estimates of *p*_0_ can actually be interpreted as the admixture proportion to which the frequency of Neanderthal alleles settled down to after the first few generations of segregation off of unlinked deleterious alleles. As a consequence, the true initial admixture proportion may have been much higher than our current estimates of *p*_0_. However, any attempt to correct for this potential bias in our estimates of *p*_0_ is likely very sensitive to assumptions about the form of selection, as we discuss below. Conversely, our estimates of the strength and density of deleterious sites (*s* and *μ*) do not strongly change when we include multiple deleterious sites or consider large windows surrounding each focal neutral site (up to 10 cM) in our inference procedure (see S2 Text for details). This is likely because much of the information about *s* and *μ* comes from the localized dip in Neanderthal ancestry close to genes, and thus these estimates are not strongly affected by the inclusion of other weakly linked deleterious alleles (the effects of which are more uniform, and mostly affect *p*_0_).

If the predicted drop in hybrid fitness is due to the accumulation of many weakly deleterious alleles in Neanderthals, as supported by our simulations, it also suggests that Neanderthals may have had a very substantial genetic load (more than 94% reduction in fitness) compared to AMH (see also [27, 28, 31]). It is tempting to conclude that this high load strongly contributed to the low population densities, and the extinction (or at least absorption), of Neanderthals when faced with competition from modern humans. However, this ignores a number of factors. First, selection against this genetic load may well have been soft, i.e. fitness is measured relative to the most fit individual in the local population, and epistasis among these many alleles may not have been multiplicative [46–48]. Therefore, Neanderthals, and potentially early-generation hybrids, may have been shielded from the predicted selective cost of their load. Second, Neanderthals may have evolved a range of compensatory adaptations to cope with this large deleterious load. Finally, Neanderthals may have had a suite of evolved adaptations and cultural practices that offered a range of fitness advantages over AMH at the cold Northern latitudes that they had long inhabited [49, 50]. These factors also mean that our estimates of the total genetic load of Neanderthals, and indeed the fitness of the early hybrids, are at best provisional. The increasing number of sequenced ancient Neanderthal and human genomes from close to the time of contact [16, 51, 52] will doubtlessly shed more light on these parameters. However, some of these questions may be fundamentally difficult to address from genomic data alone.

Whether or not the many weakly deleterious alleles in Neanderthals were a cause, or a consequence, of the low Neanderthal effective population size, they have had a profound effect on patterning levels of Neanderthal introgression in our genomes. More generally, our results suggest that differences in effective population size and nearly neutral dynamics may be an important determinant of levels of introgression across species and along the genome. Species coming into secondary contact often have different demographic histories (e.g. as is the case of *Drosophila yakuba and D. santomea [53, 54] or in Xiphophorus sister species [55]) and so the dynamics we have described may be common.*

We have here considered the case of introgression from a small population (Neanderthals) into a larger population (humans), where selection acts genome-wide against deleterious alleles introgressing. However, from the perspective of a small population with segregating or fixed deleterious alleles, introgression from a population lacking these alleles can be favoured [56]. This could be the case if the source population had a large effective size, and hence lacked a comparable load of deleterious alleles. Therefore, due to this effect, our results may also imply that Neanderthal populations would have received a substantial amount of adaptive introgression from modern humans.

## Methods

### Model

Here we describe the model for the frequency of a Neanderthal-derived allele at a neutral locus linked to a single deleterious allele. In S1 Text we extend this model to deleterious alleles at multiple linked loci. Let *S*_1_ and *N*_1_ be the introgressed (Neanderthal) alleles at the selected and linked neutral autosomal locus, respectively, and *S*_2_ and *N*_2_ the corresponding resident (human) alleles. The recombination rate between the two loci is *r*. We assume that allele *S*_1_ is deleterious in humans, such that the viability of a heterozygote human is *w* (*S*_1_*S*_2_) = 1*− s*, while the viability of an *S*_2_*S*_2_ homozygote is *w*(*S*_2_*S*_2_) = 1. We ignore homozygous carriers of allele *S*_1_, because they are expected to be very rare, and omitting them does not affect our results substantially (S1 Text). We assume that, prior to admixture, the human population was fixed for alleles *S*_2_ and *N*_2_, whereas Neanderthals were fixed for alleles *S*_1_ and *N*_1_. After a single pulse of admixture, the frequency of the introgressing haplotype *N*_1_*S*_1_ rises instantaneously from 0 to *p*_0_ in the human population (We discuss the consequences of multiple pulses in S1 Text).

In S1 Text and S2 Text we study the more generic case where both *S*_1_ and *S*_2_ are segregating in the Neanderthal population prior to admixture. Fitting this full model to data (S2 Text), we found that it resulted in estimates which implied that the deleterious allele *S*_1_ is on average fixed in Neanderthals. This was further supported by our individual-based simulations (S18 Fig), which show that in a vast majority of realisations, the deleterious allele was either at very low or very high frequency in the Neanderthals immediately prior to introgression due to the high levels of genetic drift in Neanderthals. Therefore, we focus only on the simpler model where allele *S*_1_ is fixed in Neanderthals, as described above.

The present-day expected frequency of allele *N*_1_ in modern humans can be written as

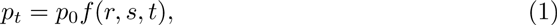

where *f* (*r, s, t*) is a function of the recombination rate *r* between the neutral and the selected site, the selection coefficient *s*, and the time *t* in generations since admixture (S1 Text).

Based on the derivations in S1 Text, we find that, for autosomes, *f* is given by

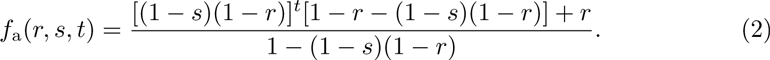

We also have developed results for a neutral locus linked to a single deleterious locus in the non-pseudo-autosomal (non-PAR) region of the X chromosome (S1 Text). As above, we also assume that the deleterious allele is fixed in Neanderthals. The non-PAR region does not recombine in males and we assume that the recombination rate in females between the two loci is *r*. In S1 Text we develop a full model allowing for sex-specific fitnesses. For simplicity, here we assume that heterozygous females and hemizygous males carrying the deleterious Neanderthal allele have relative fitness 1 − *s*. Following our results in S1 Text we obtain

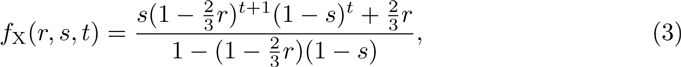

where the factors 2/3 and (1 − 2/3) reflect the fact that, on average, an X-linked allele spends these proportions of time in females and males, respectively. We also fitted models with different selection coefficients in heterozygous females and hemizygote males, but found that there was little information to separate these effects.

Our results relate to a long-standing theory on genetic barriers to gene flow [21–26], a central insight of which is that selection can act as a barrier to neutral gene flow. This effect can be modelled as a reduction of the neutral migration rate by the so-called gene flow factor [22], which is a function of the strength of selection and the genetic distance between neutral and selected loci. In a single-pulse admixture model at equilibrium, *f* is equivalent to the gene flow factor (S1 Text).

Lastly, we introduce a parameter *μ* to denote the probability that any given exonic base is affected by purifying selection. If *μ* and *s* are small, we found that considering only the nearest-neighboring selected exonic site is sufficient to describe the effect of linked selected sites in our case (but see Results and Discussion for the effect of unlinked sites under selection). That is, for small *μ*, selected sites will be so far apart from the focal neutral site ℓ that the effect of the nearest selected exonic site will dominate over the effects of all the other ones. In S1 Text we provide predictions for the present-day frequency of *N*_1_ under a model that accounts for multiple linked selected sites, both for autosomes and the X chromosome. We further assume that an exon of length *l* bases will contain the selected allele with probability *≈ μl* (for *μl* ≪ 1), and that the selected site is located in the middle of that exon. Lastly, the effects of selection at linked sites will be small if their genetic distance from the neutral site is large compared to the strength of selection (*s*). In practice, we may therefore limit the computation of Eq. (1) to exons within a window of a fixed genetic size around the neutral site. We chose windows of size 1 cM around the focal neutral site ℓ, but also explored larger windows of size 10 cM to show that our results are not strongly affected by this choice. Taken together, these assumptions greatly simplify our computations and allow us to calculate the expected present-day frequency of the Neanderthal allele at each SNP along the genome.

Specifically, consider a genomic window of size 1 cM centered around the focal neutral site ℓ, and denote the total number of exons in this window by ℐ_ℓ_. Let the length of the *i*^th^ nearest exon to the focal locus ℓ be *l_i_* base pairs. The probability that the *i*^th^exon contains the nearest selected site is then 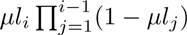, where the product term is the probability that the selected site is not in any of the *i* − 1 exons closer to ℓ than exon *i*. Conditional on the *i*^th^ exon containing the selected site, the frequency *p_t_* of *N*_1_ at locus ℓ and time *t* is computed according to Eq. (1), with *r* replaced by *r_i_*, the recombination rate between ℓ and the center of exon *i*. Then, we can write the expected frequency of the neutral Neanderthal allele at site ℓ surrounded by ℐ_*ℓ*_ exons as

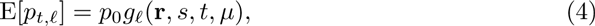

where

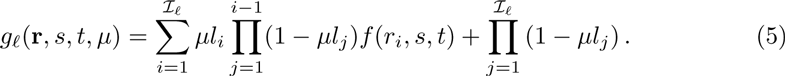

The last product term accounts for the case where none of the ℐ_*ℓ*_ exons contains a deleterious allele. Equation (5) can be applied to both autosomes and X chromosomes, with *f* as given in equations (2) and (3), respectively.

### Inference procedure

We downloaded recently published estimates of Neanderthal alleles in modern-day humans [12], as well as physical and genetic positions of polymorphic sites (SNPs) from the Reich labwebsit e. We use estimates from Sankararaman *et al.* [12] of the average marginal probability that a human individual carries a Neanderthal allele as our Neanderthal allele frequency, *p_n_*. Although *p_n_* is also an estimate, we generally refer to it as the observed frequency, in contrast to our predicted/expected frequency *p_t_*. Sankararaman *et al.* [12] performed extensive simulations to demonstrate that these calls were relatively unbiased. We performed separate analyses using estimates of *p_n_* for samples originating from Europe (EUR) and East Asia (ASN) (Table 1, [12]).

Although composed of samples from multiple populations, for simplicity we refer to EUR and ASN as two samples or populations. We downloaded a list of exons from the UCSC Genome browser. We matched positions from the GRCh37/hg19 assembly to files containing estimates of *p_n_* to calculate distances to exons. We estimated recombination rates from a genetic map by Kong *et al.* [59].

Our inference method relies on minimizing the residual sum of squared differences (RSS) between E[*p_t,ℓ_*] and *p_n,ℓ_* over all *n_l_* autosomal (or X-linked) SNPs for which [12] provided estimates. Specifically, we minimize

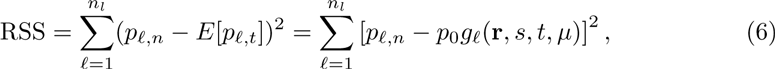

where *g_ℓ_*(**r**, *s, t,*) is calculated according to Eq. (5). For each population, we first performed a coarse search over a wide parameter space followed by a finer grid search in regions that had the smallest RSS. For each fine grid, we calculated the RSS for a total of 676 (26x26) different combinations of *s* and *μ*. We did not perform a grid search for *p*_0_. Rather, for each combination of *s* and *μ*, we analytically determined the value of *p*_0_ that minimizes the RSS as

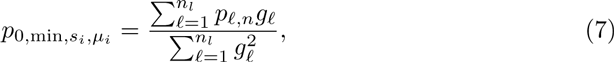

where *g_ℓ_* is given in Eq. (5) and we sum over all *n_l_* considered autosomal (X-linked) SNPs. For details, we refer to S2 Text.

We created confidence intervals by calculating 2.5 and 97.5 percentiles from 1000 bootstrapped genomes. We created these chromosome by chromosome as follows. For a given chromosome, for each non-overlapping segment of length 5 cM, and for each of 676 parameter combinations, we first calculated the denominator and the numerator of Eq. (7) using the number of SNPs in the segments instead of *n_l_*. We then resampled these segments (with replacement) to create a bootstrap chromosome of the same length as the original chromosome. Once all appropriate bootstrap chromosomes were created (chromosomes 1–22 in the autosomal case, or the X chromosome otherwise), we obtained for each bootstrap sample the combination of *p*_0_, *μ*, and *s* that minimises the RSS according to equations (6) and (7).

In S2 Text we extend our inference approach to incorporate the influence of multiple selected loci on levels of introgression (in various size windows up to 10cM in size). We also explored using a more stringent set of Neanderthal calls and using a variance-weighted sum of squares approach. All of these approaches resulted in similar estimates of *s* and *μ*, suggesting that our findings are reasonably robust to our choices.

### Individual-based simulations

To test whether selection against alleles introgressed from Neanderthals can be explained by the differences in ancient demography, we simulated the frequency trajectories of deleterious alleles in the Neanderthal and human populations, between the time of the Neanderthal–human split and the time of admixture (S3 Text). We assume that the separation time was 20,000 generations (∼ 600k years). For the distribution of selection coefficients we use those of [30]. This distribution was estimated under the assumption of no dominance [30], and we follow this assumption in our simulations. For the simulations summarized in Fig 5 we assumed an effective population size of 1000 for Neanderthals and 10,000 for humans. Our simulations are described more fully in S3 Text, where we also show versions of Fig 5 for a range of effective population sizes for Neanderthals. The timing of the out-of-Africa bottleneck in humans relative to admixture with Neanderthals is unclear. Therefore, we also explored the effect of a population bottleneck in humans (before admixture) on the accumulation of deleterious alleles (see S3 Text). We allowed the duration of this bottleneck to vary from 10 to 1000 generations. These simulations show that our findings in Fig 5 are robust to the precise details of the demography of the human populations. We acknowledge that our understanding of the human populations that initially encountered Neanderthals is scant, and they may have been small in size. However, importantly the populations that represent the ancestors of modern-day Eurasians do not appear to have had the sustained history of small effect population sizes over hundreds of thousands of years that characterize Neanderthals. Therefore, our simulations likely capture the important broad dynamics of differences in effective population size on deleterious allele load.

For each simulation run, we recorded the frequency of the deleterious allele in Neanderthals and humans immediately prior to admixture. Our simulations show that the majority of deleterious alleles that are still segregating at the end of the simulation are fixed differences (Fig 5). This matches the assumption of our approach, and agrees with the estimates we obtained. Our simulations include both ancestral variation and new mutations, but the majority of the segregating alleles at the end of the simulations represent differentially sorted ancestral polymorphisms.

Harris and Nielsen [31] independently conducted a simulation study of the accumulation of deleterious alleles in Neanderthals, and the fate of these after introgression into modern humans. Their results about the accumulation of weakly deleterious additive alleles in Neanderthals are consistent with ours. In addition, these authors also investigated the introgression dynamics with linked recessive deleterious alleles. They found that, under some circumstances, recessive deleterious alleles may actually favor introgression as a consequence of pseudo-overdominance. However, the majority of weakly selected alleles are expected to act in a close-to-additive manner, as empirical results suggest an inverse relationship between fitness effect and dominance coefficient [57, 58]. Therefore, our assumptions of additivity are appropriate for the majority of deleterious loci.

## Acknowledgements

We would like to thank Nicolas Bierne, Jeremy Berg, Vince Buffalo, Gideon Bradburd, Yaniv Brandvain, Nancy Chen, Henry Coop, Kristin Lee, Samantha Price, Alisa Sedghifar, Guy Sella, Michael Turelli, Tim Weaver, Chenling Xu, and members of the Ross-Ibarra and Schmitt labs at UC Davis for helpful feedback on the work described in this paper. We thank David Reich, Molly Schumer, and two anonymous reviewers for feedback on an earlier version of the paper. This work was supported by an Advanced Postdoc. Mobility fellowship from the Swiss National Science Foundation P300P3 154613 to SA, and by grants from the National Science Foundation under Grant No. 1353380 to John Willis and GC and the National Institute of General Medical Sciences of the National Institutes of Health under award numbers NIH RO1GM83098 and RO1GM107374 to GC.

## Supporting Information

### S1 Text

**Modeling Selection Against Introgression.** Here, we describe several models of a single pulse of admixture between Neanderthal and modern humans, and derive approximations for the present-day frequency of a neutral introgressed Neanderthal allele linked to one or multiple sites under purifying selection in humans. We then demonstrate the accuracy of these approximations by comparing them to numerically iterated recursion equations and individual-based simulations. Lastly, we consider models of single and multiple waves of continuous introgression and show that one cannot distinguish between these models and a single-pulse admixture model using the present-day frequency of introgressed alleles as the only source of information.

### S2 Text

**Inference Procedure.** Here, we introduce the last model parameter, the average probability *μ* that, at any given exonic base pair, a deleterious Neanderthal allele is segregating in the modern human population. We then discuss the details of our inference procedure and expand on our results.

### S3 Text

**Individual-based Simulations.** Here, we describe individual-based simulations to investigate whether the difference in population size between Neanderthals and modern humans can account for the selection coefficient (*s*) and the exonic density of deleterious sites (*μ*) that we estimated (main text, S2 Text).

### S1 Fig

**Approximate frequency** *p_t_* **of** *N*_1_ **as a function of the recombinational distance** *r*. Lines represent Eq. (6) for *t* = 2000 (red) and the equilibrium given in Eq. (8) (grey). Numerical iterations of the corresponding recursion equations are represented by red upward and black downward facing triangles. Other parameters are *s* = 0.0001, and *y*_0_ = 0 for all lines, and *p*_0_ = 0.04 (dotted), 0.034 (dashed) and 0.03 (full line).

### S2 Fig

**Approximate frequency** *p_t_* **of** *N*_1_ **as a function of the recombinational distance** *r*. Lines represent Eq. (6) for *t* = 2000 (red) and the equilibrium given in Eq. (8) (grey). Numerical iterations of the corresponding recursion equations are represented by red upward and black downward facing triangles. Other parameters are *s* = 0.0004, and *y*_0_ = 0 for all lines, and *p*_0_ = 0.04 (dotted), 0.034 (dashed) and 0.03 (full line).

### S3 Fig

**Approximate frequency** *p_t_* **of** *N*_1_ **as a function of the recombinational distance** *r* **for the X chromosome.** Lines represent Eq. (12) for *t* = 2000 (red) and the equilibrium from Eq. (13) (grey). Numerical iterations of the corresponding recursion equations are represented by red upward and black downward facing triangles. Other parameters are *s_f_* = *s_m_* = 0.0001, and *y*_*X*,0_ = 0 for all lines, and *p*_0_ = 0.04 (dotted), 0.034 (dashed) and 0.03 (full line).

### S4 Fig

**Approximate frequency** *p_t_* **of** *N*_1_ **as a function of the recombinational distance** *r* **for the X chromosome.** Lines represent Eq. (12) for *t* = 2000 (red) and the equilibrium from Eq. (13) (grey). Numerical iterations of the corresponding recursion equations are represented by red upward and black downward facing triangles. Other parameters are *s_f_* = *s_m_* = 0.0004, and *y_X_*,_0_ = 0 for all lines, and *p*_0_ = 0.04 (dotted), 0.034 (dashed) and 0.03 (full line).

### S5 Fig

**Comparison of the mean frequency of** *N*_1_ **obtained from individual-based simulations to the theoretical prediction from Eq. (6).** The figure shows 676 circles representing different combinations of *r* (recombination rate) and *s* (selection coefficient). Values of *r* range from 1 *×* 10^−5^ (red circle border) to 1 *×* 10^−2^ (black border), *s* ranges from 1 *×* 10^−5^ (yellow circle area) to 4 *×* 10^−4^ (light blue area). For each parameter combination, the mean frequency of *N*_1_ after *t* = 2000 generations was calculated across 1000 independent runs. Grey lines represent approximate 95% confidence intervals for simulation results (mean *±* 1.96 *×* standard error), and a black line with slope 1 is shown for reference.

### S6 Fig

**Accuracy of approximation to the frequency of a neutral allele** *N*_1_ **linked to multiple autosomal loci under purifying selection.** Curves show *p_∞,I J_* from Eq. (15) for various recombination distances between the focal neutral locus N and the two loci under selection, A and B. Upward and downward facing triangles give values obtained after iterating deterministic recursions over *t* = 2000 generations and until the equilibrium is reached, respectively. A: The neutral locus is flanked by one locus under selection on each side, and recursions followed Eq. (17). B: The neutral locus is flanked by two selected loci on one side and recursions followed Eq. (18). A, B: Selection coefficients against introgressed deleterious mutations at locus A and B are *a* = 0.0002 and *b* = 0.0004, respectively. The initial frequency of *N*_1_ is *p*_0_ = 0.04.

### S7 Fig

**Accuracy of approximation to the frequency of a neutral allele** *N*_1_ **linked to multiple X-chromosomal loci under purifying selection.** Curves show *p_X,∞,I J_* from Eq. (21) for various recombination distances between the focal neutral locus N and the two loci under selection, A and B. Upward and downward facing triangles give values obtained after iterating Eq. (24) over *t* = 2000 generations and until the equilibrium is reached, respectively. A, B: The neutral locus is flanked by one locus under selection on each side. C, D: The neutral locus is flanked by two loci under selection on one side. A, C: Selection coefficients against introgressed deleterious mutations at locus A and B in females (males) are *a_f_* = 0.0001 (*a_m_* = 0.0003) and *b_f_* = 0.0002 (*b_m_* = 0.0006), respectively. B, D: Selection coefficients are identical in the two sexes; *a_f_* = *a_m_* = 0.0001 and *b_f_* = *b_m_* = 0.0002. In all panels, the initial frequency of *N*_1_ is *p*_*X*,0_ = 0.04.

### S8 Fig

**Mapping models with one (red line) and two (blue line) waves of introgression to a single-pulse model.** By changing time in the single-pulse model (dashed and dotted black lines) as described in S1 Text, we can recover present-day haplotype frequencies generated by the wave models. Parameters are *r* = 10^−4^, *s* = 5 *×* 10^−4^, *x*_0_ = 0.04, and *y*_0_ = 0.001. The duration of admixture in the single-wave model is *τ* = 500. Additional parameters for the dual-wave model are *τ*_1_ = 75, *τ*_2_ = 1075, *τ*_3_ = 1500. The solid black line represents a single-pulse model without change of time.

### S9 Fig

**The scaled RSS surface** (RSS_min_ *−* RSS) **for different** *s* **and** *μ* **values for EUR and ASN autosomal chromosomes under the single-locus equilibrium model** (*t* = *∞*). Each value of the RSS is minimized over *p*_0_, making this a profile RSS surface. Regions shaded in orange represent parameter values of higher RSS.

### S10 Fig

**The scaled RSS surface** (RSS_min_ *−* RSS) **for different** *s* **and** *μ* **values for EUR and ASN autosomal chromosomes under the single-locus model for** *t* = 2000. Each value of the RSS is minimized over *p*_0_, making this a profile RSS surface. Regions shaded in orange represent parameter values of higher RSS. Black circles show bootstrap results of 1000 block bootstrap reestimates, with darker circles corresponding to more common bootstrap estimates.

### S11 Fig

**The scaled RSS surface** (RSS_min_ *−* RSS) **for different** *s* **and** *μ* **values for EUR and ASN autosomal chromosomes under a multi-locus equilibrium model** (*t* = *∞*). Each value of the RSS is minimized over *p*_0_, making this a profile RSS surface. Regions shaded in orange represent parameter values of higher RSS.

### S12 Fig

**The scaled RSS surface** (RSS_min_ *−* RSS) **for different** *s* **and** *μ* **values for the X chromosome in the ASN population under a single-locus model for** *t* = 2000 **and assuming equal strength of selection in males and females.** Each value of the RSS is minimized over *p*_0_, making this a profile RSS surface. Regions shaded in orange represent parameter values of higher RSS. Black circles show bootstrap results of 1000 block bootstrap reestimates, with darker circles corresponding to more common bootstrap estimates.

### S13 Fig

**The scaled RSS surface** (RSS_min_ *−* RSS) **for different** *s* **and** *μ* **values for the X chromosome in the ASN population for a single-locus model for** *t* = 2000 **and assuming equal strength of selection in males and females.** Each value of the RSS is minimized over *p*_0_, making this a profile RSS surface. Regions shaded in orange represent parameter values of higher RSS. Black circles show bootstrap results of 1000 block bootstrap reestimates, with darker circles corresponding to more common bootstrap estimates.

### S14 Fig

**The scaled RSS surface** (RSS_min_ *−* RSS) **for the X chromosomes as a function of the initial admixture proportion** *p*_0_. Results are shown for a model where only the nearest-neighboring exonic site under selection is considered, and for *t* = 2000 generations after Neanderthals split from the EUR (grey) and ASN (pink) populations. Dots and horizontal lines show the value of *p*_0_ that minimizes the RSS and the respective 95% block-bootstrap confidence intervals. Each value of the RSS is evaluated at the values of the selection coefficient (*s*) and exonic density of selection (*μ*) given in Table.

### S15 Fig

**Fit between our estimates of** *p_t_* **for bins of different exon density.** Genomic regions with low exonic density (low exonic density rank) contain higher average Neanderthal allele frequency in both in Europeans (grey circle) and Asians (pink circle), a pattern recreated in our model. Dashed lines represent the 95% block bootstrap confidence intervals. The length of segments used to create the bins is 2 cM.

### S16 Fig

**Fit between our estimates of** *p_t_* **for bins of different exon density.** Genomic regions with low exonic density (low exonic density rank) contain higher average Neanderthal allele frequency in both in Europeans (grey circle) and Asians (pink circle), a pattern recreated in our model. Dashed lines represent the 95% block bootstrap confidence intervals. The length of segments used to create the bins is 1.5 cM.

### S17 Fig

**Fit between our estimates of** *p_t_* **for bins of different exon density.** Genomic regions with low exonic density (low exonic density rank) contain higher average Neanderthal allele frequency in both in Europeans (grey circle) and Asians (pink circle), a pattern recreated in our model. Dashed lines represent the 95% block bootstrap confidence intervals. The length of segments used to create the bins is 0.5 cM. There are 9 bins, rather than 10 bins, in this figure because there are many 0.5 cM bins with zero exonic sites. Therefore, we collapsed our results together into a smaller number of bins.

### S18 Fig

**The scaled RSS surface** (RSS_min_ *−* RSS) **for different values of** *s* **and** *μ* **for EUR and ASN autosomes under a multi-locus equilibrium model** (*t* = *∞*). This surface is constructed using windows of 10 cM, but otherwise analogous to. Each value of the RSS is minimized over *p*_0_, which makes this a profile RSS surface. Regions shaded in orange represent parameter values of higher RSS.

### S19 Fig

**The scaled RSS surfaces** (RSS_min_ *−* RSS) **for different values of** *s* **and** *μ* **for the X chromosome under a multi-locus equilibrium model** (*t* = *∞*). This surface is constructed using windows of 10 cM. Each value of the RSS is minimized over *p*_0_, which makes this a profile RSS surface. Regions shaded in orange represent parameter values of higher RSS.

### S20 Fig

**The scaled RSS surfaces** (RSS_min_ *−* RSS) **for different** *s* **and** *μ* **values for EUR and ASN autosomes under a single-locus model** (*t* = 2000). This surface is constructed using the fraction of EUR and ASN alleles at each site with confident Neanderthal calls (a marginal probability of *>* 90%). Each value of the RSS is minimized over *p*_0_, which makes this a profile RSS surface. Regions shaded in orange represent parameter values of higher RSS. The window size 1 cM.

### S21 Fig

**Comparison of the variance and the mean frequency of** *N*_1_ **obtained from individual-based simulations**. The figure shows 676 circles representing different combinations of *r* (recombination rate) and *s* (selection coefficient). Values of *r* range from 1 *×* 10^−5^ (red circle border) to 1 *×* 10^−2^ (black border), *s* ranges from 1 *×* 10^−5^ (yellow circle area) to 4 *×* 10^−4^ (light blue area). For each parameter combination, the mean and variance of the frequency of *N*_1_ after *t* = 2000 generations was calculated across 1000 independent runs.

### S22 Fig

**The scaled weighted RSS surface** (RSS_min_ *−* RSS) **for different** *s* **and** *μ* **values for EUR and ASN autosomal chromosomes under the single-locus model for** *t* = 2000. Each value of the RSS is minimized over *p*_0_, which makes this a profile RSS surface. The window size 1 cM.

### S23 Fig

**Simulations showing that the Neanderthal population is predicted to harbor an excess of weakly deleterious fixed alleles compared to humans.** In panel A we show a 2d histogram of the difference in allele frequency between the Neanderthal to human and the deleterious selection coefficient over all sites in our simulations. In panel B we show the fraction of sites in the simulations where there is a human- or Neanderthal-specific fixed differences binned by selection coefficient. In panel B we show the fraction of sites in the simulations where there is a human- or Neanderthal-specific fixed differences binned by selection coefficient. In B we mark with dotted lines the nearly-neutral selection coefficient boundary for Neanderthal (right) and Human (left) populations and with solid lines our 95% CI of *s* for ASN (the larger of the two CI). Note that monomorphic sites are not shown, but are included in the denominator of the fraction of sites. In these simulations *N_n_* = 500 and *u* = 10^−8^.

### S24 Fig

**The same as S18 Fig except that** *N_n_* = 1000.

### S25 Fig

**The same as S18 Fig, except that** *N_n_* = 2000.

### S26 Fig

**As in S23 Fig, but with a bottleneck in the human population of length** *T_b_* = 10 **generations prior to admixture with Neanderthals.** The long-term effective size of the human population prior to the bottleneck was set to *N_h_* = 14,400, and the effective size during the bottleneck to 1861 (see S3 Text for details).

### S27 Fig

**As in S26 Fig, but with a bottleneck duration of** *T_b_* = 100 **generations**

### S28 Fig

**As in S26 Fig, but with a bottleneck duration of** *T_b_* = 1000 **generations**

### S1 Table

**Minimum RSS parameters for** *μ*, *s* **and** *p*_0_ **for different models described in S1 Text**. Fig 1 in the main text shows an example of *E*[*p_t_*] for single locus model, *t* = 2000, for part of chromosome 1.

### S2 Table

**The 95% bootstrap confidence intervals for** *μ*, *s*, **and** *p*_0_ **for different models.**

### S3 Table

**Correlation between the estimated and the observed mean Neanderthal allele frequency for bins created using segments of different sizes.**

## References

1. Noonan JP, Coop C, Kudaravalli S, Smith D, Krause J, Alessi J, et al. Sequencing and analysis of neanderthal genomic dna. Science. 2006;314(5802): 1113–1118. doi: 10.1126/science.1131412

2. Green RE, Krause J, Briggs AW, Maricic T, Stenzel U, Kircher M, et al. A Draft Sequence of the Neandertal Genome. Science. 2010;328(5979): 710–722. doi: 10.1126/science.1188021

3. Reich D, Green RE, Kircher M, Krause J, Patterson N, Durand EY, et al. Genetic history of an archaic hominin group from Denisova Cave in Siberia. Nature. 2010;468(7327): 1053–1060.

4. Meyer M, Kircher M, Gansauge MT, Li H, Racimo F, Mallick S, et al. A High-Coverage Genome Sequence from an Archaic Denisovan Individual. Science. 2012;338(6104): 222–226. doi: 10.1126/science.1224344

5. Prüefer K, Racimo F, Patterson N, Jay F, Sankararaman S, Sawyer S, et al. The complete genome sequence of a Neanderthal from the Altai Mountains. Nature. 2014;505(7481): 43–49. doi: 10.1038/nature12886

6. Sankararaman S, Patterson N, Li H, Pääbo S, Reich D. The date of interbreeding between Neandertals and modern humans. PLoS Genet. 2012;8(10): e1002947. doi: 10.1371/journal.pgen.1002947

7. Wall JD, Yang MA, Jay F, Kim SK, Durand EY, Stevison LS, et al. Higher levels of neanderthal ancestry in East Asians than in Europeans. Genetics. 2013;194(1): 199–209.

8. Vernot B, Akey JM. Resurrecting Surviving Neandertal Lineages from Modern Human Genomes. Science. 2014;343(6174): 1017–1021. doi: 10.1126/science.1245938

9. Vernot B, Akey JM. Complex history of admixture between modern humans and Neandertals. Am J Hum Genet. 2015;96(3): 448–453.

10. Kim BY, Lohmueller KE. Selection and reduced population size cannot explain higher amounts of Neandertal ancestry in East Asian than in European human populations. Am J Hum Genet. 2015;96(3):454–461.

11. Khrameeva EE, Bozek E, He L, Yan Z, Jiang X, Wei Y, et al. Neanderthal ancestry drives evolution of lipid catabolism in contemporary Europeans. Nat Commun. 2014; doi: 10.1038/ncomms4584.

12. Sankararaman S, Mallick S, Dannemann M, Prüefer K, Kelso J, Paeaebo S, et al. The genomic landscape of Neanderthal ancestry in present-day humans. Nature. 2014;507(7492): 354–357. doi:10.1038/nature12961

13. Racimo F, Sankararaman S, Nielsen R, Huerta-Sanchez E. Evidence for archaic adaptive introgression in humans. Nat Rev Genet. 2015;16(6): 359–371. doi:10.1038/nrg3936

14. Serre D, Langaney A, Chech M, Teschler-Nicola M, Paunovic M, Mennecier P. No evidence of Neandertal mtDNA contribution to early modern humans. PLoS Biol. 2004;2(3): e57.

15. Currat M, Excoffier L. Modern humans did not admix with neanderthals during their range expansion into europe. PLoS Biol, 2(12):e421, 2004.

16. Fu Q, Posth C, Hajdinjak M, Petr M, Mallick S et al. The genetic history of Ice Age Europe. Nature. 2016.

17. Sankararaman S, Mallick S, Patterson N, Reich D. The Combined Landscape of Denisovan and Neanderthal Ancestry in Present-Day Humans. Current Biology. 2016.

18. Vernot B, Tucci S, Kelso J, Schraiber JG, Wolf AB, Gittelman RM et al. Excavating Neandertal and Denisovan DNA from the genomes of Melanesian individuals. Science. 2016; 352 (6282): 235–239.

19. Currat M, Excoffier L. Strong reproductive isolation between humans and Neanderthals inferred from observed patterns of introgression. Proc Natl Acad Sci USA. 2011;108(37): 15129–15134.

20. Gibbons A. Neandertals and moderns made imperfect mates. Science. 2014;343(6170): 471–472.

21. Petry D. The effect on neutral gene flow of selection at a linked locus. Theor Popul Biol. 1983;23(3): 300–313. doi: 10.1016/0040-5809(83)90020-5.

22. Bengtsson, BO. The flow of genes through a genetic barrier. In: Greenwood JJ, Harvey PH, Slatkin M, editors. Evolution Essays in honour of John Maynard Smith. Cambridge: Cambridge University Press; 1985. pp. 31–42.

23. Barton NH, Bengtsson BO. The barrier to genetic exchange between hybridizing populations. Heredity. 1986;57(3): 357–376. doi: 10.1038/hdy.1986.135.

24. Charlesworth B, Nordborg M, Charlesworth D. The effects of local selection, balanced polymorphism and background selection on equilibrium patterns of genetic diversity in subdivided populations. Genet Res. 1997;70(2): 155–174

25. Gavrilets S, Hybrid zones with dobzhansky-type epistatic selection. Evolution. 1997;51(4): 1027–1035.

26. Gavrilets S, Cruzan MB. Neutral gene flow across single locus clines. Evolution. 1998;52(5): 1277–1284.

27. Do R, Balick D, Li H, Adzhubei I, Sunyaev S, Reich D. No evidence that selection has been less effective at removing deleterious mutations in Europeans than in Africans. Nat Genet. 2015;47(2): 126–131.

28. Castellano S, Parra G, Sánchez-Quinto FA, Racimo F, Kuhlwilm M, Kircher M, et al. Patterns of coding variation in the complete exomes of three neandertals. Proc Natl Acad Sci USA. 2014;111(18): 6666–6671.

29. Lin YL, Pavlidis P, Karakoc E, Ajay J, Gokcumen O. The evolution and functional impact of human deletion variants shared with archaic hominin genomes. Mol Biol Evol. 2015;32(4): 1008–1019.

30. Boyko AR, Williamson SH, Indap AR, Degenhardt JD, Hernandez RD, Lohmueller KE, et al. Assessing the evolutionary impact of amino acid mutations in the human genome. PLoS Genet. 2008;4: e1000083 doi:10.1371/journal.pgen.1000083

31. Harris K, Nielsen R. The Genetic Cost of Neanderthal Introgression. Genetics. 2016.

32. Charlesworth B, Coyne JA, Barton NH. The relative rates of evolution of sex chromosomes and autosomes. Amer Nat. 1987;130(1): 113–146.

33. Vicoso B, Charlesworth B. Evolution on the x chromosome: unusual patterns and processes. Nat Rev Genet. 2006;7(8): 645–653.

34. Meisel RP, Connallon T. The faster-X effect: integrating theory and data. Trends Genet. 2013;29(9): 537–544.

35. Wiehe TH, Stephan W. Analysis of a genetic hitchhiking model, and its application to DNA polymorphism data from Drosophila melanogaster. Mol Biol Evol. 1993;10(4): 842–854.

36. McVicker G, Gordon D, Davis C, Green P. Widespread genomic signatures of natural selection in hominid evolution. PLoS Genet. 2006;5: e1000471.

37. Sattath S, Elyashiv E, Kolodny O, Rinott Y, Sella G. Pervasive adaptive protein evolution apparent in diversity patterns around amino acid substitutions in Drosophila simulans. PLoS Genet. 2014;7: e1001302.

38. Elyashiv E, Sattath S, Hu TT, Strustovsky A, McVicker G, Andolfatto P, et al. A genomic map of the effects of linked selection in drosophila. 2014. arXiv preprint arXiv:1408.5461.

39. Rogers RL. Chromosomal Rearrangements as Barriers to Genetic Homogenization between Archaic and Modern Humans. Mol Biol Evol. 2015. in press.

40. Fitzpatrick BM. Rates of evolution of hybrid inviability in birds and mammals. Evolution. 2004;58(8) 1865–1870.

41. Curnoe D, Thorne A, Coate JA. Timing and tempo of primate speciation. J Evol Biol. 2006;19(1): 59–65.

42. Wang RJ White MA Payseur BA. The Pace of Hybrid Incompatibility Evolution in House Mice. Genetics. 2015; 201 (1): 229–242.

43. Orr HA. The population genetics of speciation: the evolution of hybrid incompatibilities. Genetics. 1995;139(4): 1805–1813.

44. Orr HA and Turelli M. The evolution of postzygotic isolation: accumulating Dobzhansky–Muller incompatibilities. Evolution. 2001;55(6): 1085–1094.

45. Dutheil JY, Munch K, Nam K, Mailund T, Schierup MH. Strong Selective Sweeps on the X Chromosome in the Human–Chimpanzee Ancestor Explain Its Low Divergence. PLoS Genet. 2015;11(8): e1005451.

46. Wallace B. Hard and soft selection revisited. Evolution, 1975;29(3): 465–473.

47. Kondrashov AS. Contamination of the genome by very slightly deleterious mutations: why have we not died 100 times over? J Theor Biol. 1995;175(4): 583–594.

48. Charlesworth B. Why we are not dead one hundred times over. Evolution 2013;67(11): 3354–3361.

49. Weaver TD. Out of Africa: modern human origins special feature: the meaning of neandertal skeletal morphology. Proc Natl Acad Sci USA 2009;106(38): 16028–16033.

50. Churchill SE. Thin on the ground: Neandertal biology, archeology and ecology. 1st ed. John Wiley & Sons; 2014.

51. Fu Q, Li H, Moorjani P, Jay F, Slepchenko SM, Bondarev AA, et al. Genome sequence of a 45,000- year-old modern human from western Siberia. Nature. 2014;514(7523): 445–449.

52. Fu Q, Hajdinjak H, Moldovan OT, Constantin S, Mallick S, Skoglund P et al. An early modern human from Romania with a recent Neanderthal ancestor. Nature. 2015;524(7564): 216–219.

53. Llopart A, Lachaise D, Coyne JA. Multilocus analysis of introgression between two sympatric sister species of *Drosophila: Drosophila yakuba* and *D. santomea*. Genetics. 2005; 171 (1): 197–210.

54. Bachtrog D, Thornton K, Clark A, Andolfatto P. Extensive introgression of mitochondrial DNA relative to nuclear genes in the *Drosophila yakuba* species group. Evolution. 2006; 60 (2): 292–302

55. Schumer M, Cui R, Powell DL, Rosenthal GG, Andolfatto P. Ancient hybridization and genomic stabilization in a swordtail fish. Mol Ecol. 2016.

56. Bierne N, Lenormand T, Bonhomme F, David P. Deleterious mutations in a hybrid zone: can mutational load decrease the barrier to gene flow? Genet Res. 2002; 80 (03): 197–204

57. Phadnis N, Fry D. Widespread Correlations Between Dominance and Homozygous Effects of Mutations: Implications for Theories of Dominance. Genetics. 2005; 171 (1): 385–392

58. Agrawal AF, Whitlock MC. Inferences About the Distribution of Dominance Drawn From Yeast Gene Knockout Data. Genetics. 2011; 187 (2): 553–566.

59. Kong A, Thorleifsson G, Gudbjartsson DF, Masson G, Sigurdsson A et al. Fine-scale recombination rate differences between sexes, populations and individuals. Nature. 2010; 467 (7319): 1099–1103.

